# Posterior simulation-based calibration tests of phylogenetic dating methods

**DOI:** 10.64898/2026.04.14.718426

**Authors:** Benedict King

## Abstract

Simulation-based calibration (SBC) checking is a method to ensure that the inference machinery for a Bayesian statistical analysis is functioning in a correct and unbiased manner. Typically, SBC begins with sampling parameter values from the model priors (prior SBC). However, it has been shown that prior SBC can miss problems when these manifest only in certain regions of parameter space. In phylogenetics, this is relevant not only because of the vastness of tree and parameter space, but also because many phylogenetic analyses involve some degree of model misspecification. Posterior SBC is a recently developed method for checking that the inference algorithms function correctly for a given empirical dataset. Here I use posterior SBC to test the implementation of phylogenetic dating methods in the inference software BEAST 2. I test both the tip-dated approach, employing an Indo-European vocabulary dataset, and the node-dated approach, employing a molecular rRNA dataset of Tabanidae (horseflies). In both cases, posterior SBC tests indicate good calibration. Despite this, posterior predictive datasets simulated from the posterior distribution provided no further increase in the precision of node age estimates compared to the original posterior, a result consistent with previous literature showing fundamental theoretical limits to the identifiability of node ages. Nevertheless, these results suggest that phylogenetic dating methods in BEAST 2 are not biased by problems with the inference machinery, thereby increasing confidence in results obtained using these methods.

## Introduction

If we are to trust the results of a Bayesian phylogenetic analysis, it is important that the correctness of the computation is verified (Mendes *et al*. 2025). One criterion for validating inference machinery is “calibration”, such that, for example, predictions with 90% confidence are true 90% of the time (Dawid 1982). A Bayesian analysis, provided it is correctly implemented, should be well-calibrated when draws from the model prior are analysed. Simulation-based calibration (SBC) checking (Talts *et al*. 2018), is one method for validating the inference algorithm.

SBC exploits the self-consistency of Bayesian inference to check that an analysis is well-calibrated. Considering the parameters θ and data y, Cook *et al*. (2006) showed that a draw of parameter values from the prior distribution

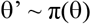

is also a valid draw from the posterior distribution

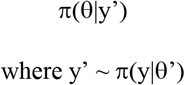

Therefore, any draw θ’’ from the posterior distribution, conditional on the data y’, should come from the same distribution as the draw from the prior distribution θ’ which generated the data y’. If θ’’ and θ’ come from the same distribution, the rank of θ’ within a set of draws from the posterior θ’’_1_, …, θ’’_s_ is uniformly distributed. In other words, the Probability Integral Transform (PIT) values, given by:

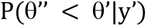

Should be drawn from a uniform distribution.

SBC proceeds in three steps. In the first step, *n* draws of the parameters are made from their prior distributions. In the second step, *n* datasets are simulated using these parameter draws. Finally, the inference algorithm is implemented to produce *n* posterior distributions, each consisting of *s* samples. For each parameter, the empirical PIT score of each of the prior draws within its respective posterior distribution is calculated to produce *n* PIT scores. If the inference algorithm is correctly implemented, these PIT scores should follow a uniform distribution. Uniformity can be visualised using histograms or an empirical cumulative distribution function (ECDF) plot (Säilynoja *et al*. 2022).

The standard form of SBC, hereafter prior SBC, checks that the algorithm performs well under parameter values sampled from the prior. However, it has been shown that even when an implementation passes prior SBC checks, problems can manifest in certain regions of parameter space (Säilynoja *et al*. 2026). For this reason, posterior SBC was developed as a way to check the validity of algorithms conditional on the data (Säilynoja *et al*. 2026).

Posterior SBC utilises the sequential nature of Bayesian inference to perform SBC in the region of parameter space occupied by the posterior. If the posterior is considered as the new prior, draws from the posterior predictive distribution can be considered as draws from this new prior. A draw from the posterior distribution θ’ ∼ π(θ|y’) is therefore also a valid draw from the augmented posterior θ’’ ∼ π(θ|y’,y’’) conditional on the predictive draw y’’ ∼ π(y|θ’). In other words, parameter draws from the posterior should come from the same distribution as draws from the augmented posterior, conditional on data from the posterior predictive distribution.

Posterior SBC follows three steps. First, *n* sets of parameters are drawn from the posterior distribution, usually by running MCMC with the empirical data. Then, *n* posterior predictive datasets are simulated using the sampled parameter values. In step three, *s* samples are drawn from the augmented posterior distribution for each of the *n* replicates, by running MCMC with both the original empirical data and a posterior predictive dataset. PIT scores of the original posterior draws, calculated as:

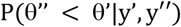

Should follow a uniform distribution when the inference algorithm is correctly implemented.

Posterior SBC is an especially important method for phylogenetics, in which the vastness of phylogenetic tree space and the complexity of phylogenetic models mean that the posterior likely occupies a region of parameter space rarely if ever sampled in the prior. Model misspecification presents a further challenge to testing modelling implementations. Posterior predictive simulations have not been routinely performed in phylogenetic analyses, but where they have some degree of model misspecification is often demonstrated (Drummond and Suchard 2009; Duchêne *et al*. 2015; Duchêne *et al*. 2019). Since model misspecification is probably widespread in phylogenetics, and given the complex nature of the analyses probably somewhat unavoidable, checking that the inference algorithms work for real empirical datasets is even more important. Posterior SBC checks the inference algorithm works even in the presence of model misspecification (Säilynoja *et al*. 2026).

Here I perform posterior SBC for two empirical phylogenetic datasets to assess the validity of phylogenetic dating methods in the software BEAST 2 (Bouckaert *et al*. 2019). The analyses cover the two main methods for calibrating the phylogenetic clock: tip-dating and node-dating. The datasets also come from disparate sources: the tip-dated analysis an Indo-European language dataset whereas the node data analyses use a molecular (rRNA) dataset.

## Methods

### Tip-dated Indo-European Language dataset

I selected a subset of the Indo-European cognate relationships database (Anderson *et al*. 2025) for analysis. I selected all meanings with 25–34 cognate sets for a total of 1336 cognate sets in 46 meanings. The analysis used a covarion substitution model (Tuffley and Steel 1998; Penny *et al*. 2001), the optimised relaxed clock (Drummond *et al*. 2006; Zhang and Drummond 2020; Douglas *et al*. 2021) and birth-death skyline tree prior (Stadler *et al*. 2013) with sampled ancestors (Gavryushkina 2014). Since this study is specifically designed to test the dating aspect of phylogenetic analysis and not tree topology, I calculated the CCD0 summary tree (Berling *et al*. 2025) from the tree sample from Heggarty *et al*. (2023), and fixed this topology during all analyses. Fixing the topology facilitates direct comparison of estimated and true node ages across the entire tree. The free parameters of the model were drawn from the following distributions:

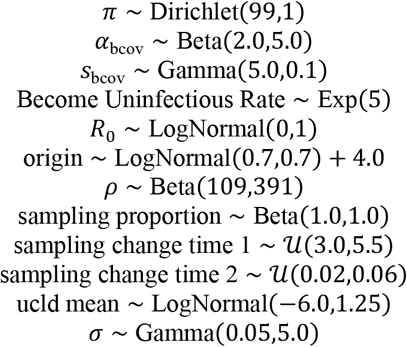

The age of each extinct language was also drawn from a normal or lognormal distribution, following Heggarty *et al*. (2023).

I drew samples from the prior by running MCMC for 20 million generations without data. For the posterior, I ran the analysis with data for three independent chains, each of 20 million generations. Convergence of the runs was confirmed in Tracer (Rambaut *et al*. 2018).

To assess model misspecification of the tree prior, I performed posterior predictive simulations for the tree model using MCMC. For each of 1000 samples from the posterior, tree model parameters taken from the posterior were fixed in the corresponding xml file and the chain run without data for 100,000 generations (sufficient to ensure burn-in of the MCMC chain), after which a single tree was logged. The resulting 1000 posterior predictive trees were combined for further analysis. The BEAST 2 package treestat2 was used to calculate statistics for both the posterior and posterior predictive trees, including external:internal branch length ratio, number of sampled ancestors, tree length and tree height and gamma (Pybus and Harvey 2000).

I performed standard (prior) SBC using 100 trees from the sample generated by MCMC without data. I simulated cognate sets on the resulting trees using the sampled parameter values of the substitution and clock models. The sampled branch rates were used directly, as opposed to resampling branch rates from the clock rate and standard deviation. The resulting simulated data was filtered so that fast and slow forms of the present and absent states were collapsed into binary present/absent, and all-absent cognates were removed. The first 1336 of the remaining simulated cognate sets were then reanalysed using the same model set-up used to generate the data. Effective sample size (ESS) scores were calculated using the R package sns (Mahani *et al*. 2016) for each free parameter in each simulation replicate. ESS scores were overwhelmingly above 200, but due to the very large number of runs being performed, lower ESS scores in a very small proportion of parameter estimates were tolerated (Table 1).

**Table 1.**
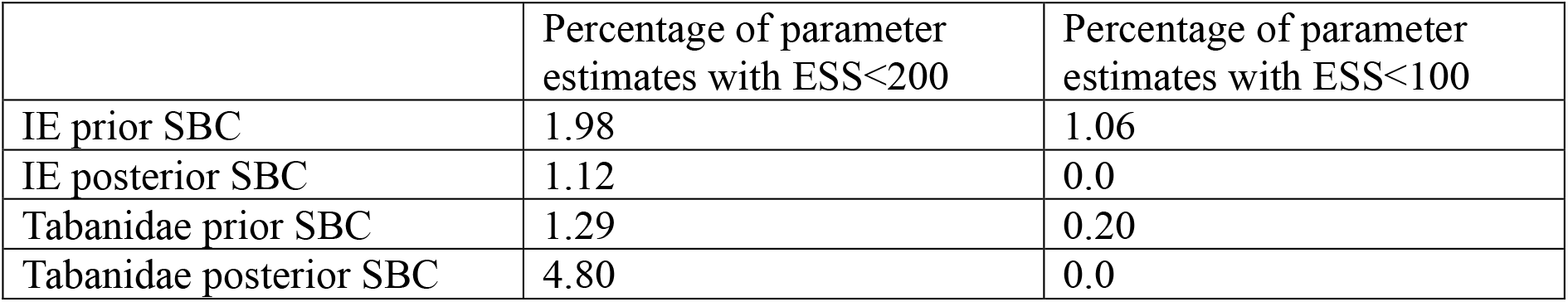
Convergence diagnostics for the SBC experiments. The number of parameter estimates in each experiment is equal to the number of parameters multiplied by the number of simulation replicates.

Simulations for posterior SBC were performed exactly as for prior SBC, but using 100 trees from the posterior sample, generated by MCMC including data. The simulated datasets were reanalysed using a two-partition set-up. One partition contained the original empirical data, and the other partition contained one of the posterior predictive datasets generated through simulation. To assess to what extent results from posterior SBC were due to properties of the empirical data, I also performed posterior SBC on one of the prior predictive datasets generated for prior SBC. In other words, following the same steps as for posterior SBC, but with a simulated dataset drawn from the prior predictive distribution in place of the empirical data.

Results of both prior and posterior SBC were checked for coverage and uniformity of PIT values. Empirical Cumulative Distribution Function (ECDF) difference plots were produced using the R package bayesplot (Gabry *et al*. 2019). ECDF difference plots show the deviation of the observed cumulative distribution of PIT scores from the expectation under uniformity (Säilynoja *et al*. 2022).

### Node-Dated Tabanidae dataset

A dataset of Tabanidae (horseflies) was selected for testing (Morita *et al*. 2016). The analysis was performed using the non-stem 28s sites from the data (1174 sites). I applied a Yule tree prior, the HKY substitution model (Hasegawa *et al*. 1985) and the uncorrelated relaxed clock. The free parameters were drawn from the following prior distributions:

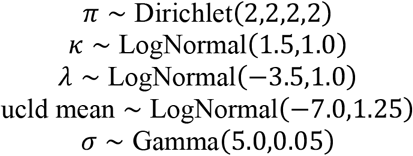

In addition, calibrations were applied to three nodes. These were applied as lognormal distributions (mean 0.5, sd 0.5) with offsets of 44 Ma (Bouvieromyiini), 112 Ma (Tabanidae) and 130 Ma (Tabanidae + Athericidae). Prior and Posterior SBC were performed as described above. No filtering and processing step was necessary for the molecular data; alignments of 1174 sites were simulated and used directly for reanalysis.

Posterior predictive simulations were again performed using MCMC, as above. The node calibration densities were included as part of the model, as opposed to simulating a pure yule process. Likewise, prior and posterior SBC followed the same steps outlined above.

## Results

### Tip-dated Indo-European language dataset

The majority of parameter estimates in the posterior distribution depart clearly from their effective prior distributions (Figure 1). This confirms that the subset of data selected was sufficiently large for posterior SBC to be a true test of the posterior rather than a recapitulation of prior SBC, but the smaller dataset size remains computationally tractable for posterior SBC. Misspecification of the tree model is demonstrated by separation of the posterior predictive and posterior trees in the tree length and sampled ancestor count metrics (Figure 2).

**Figure 1.**
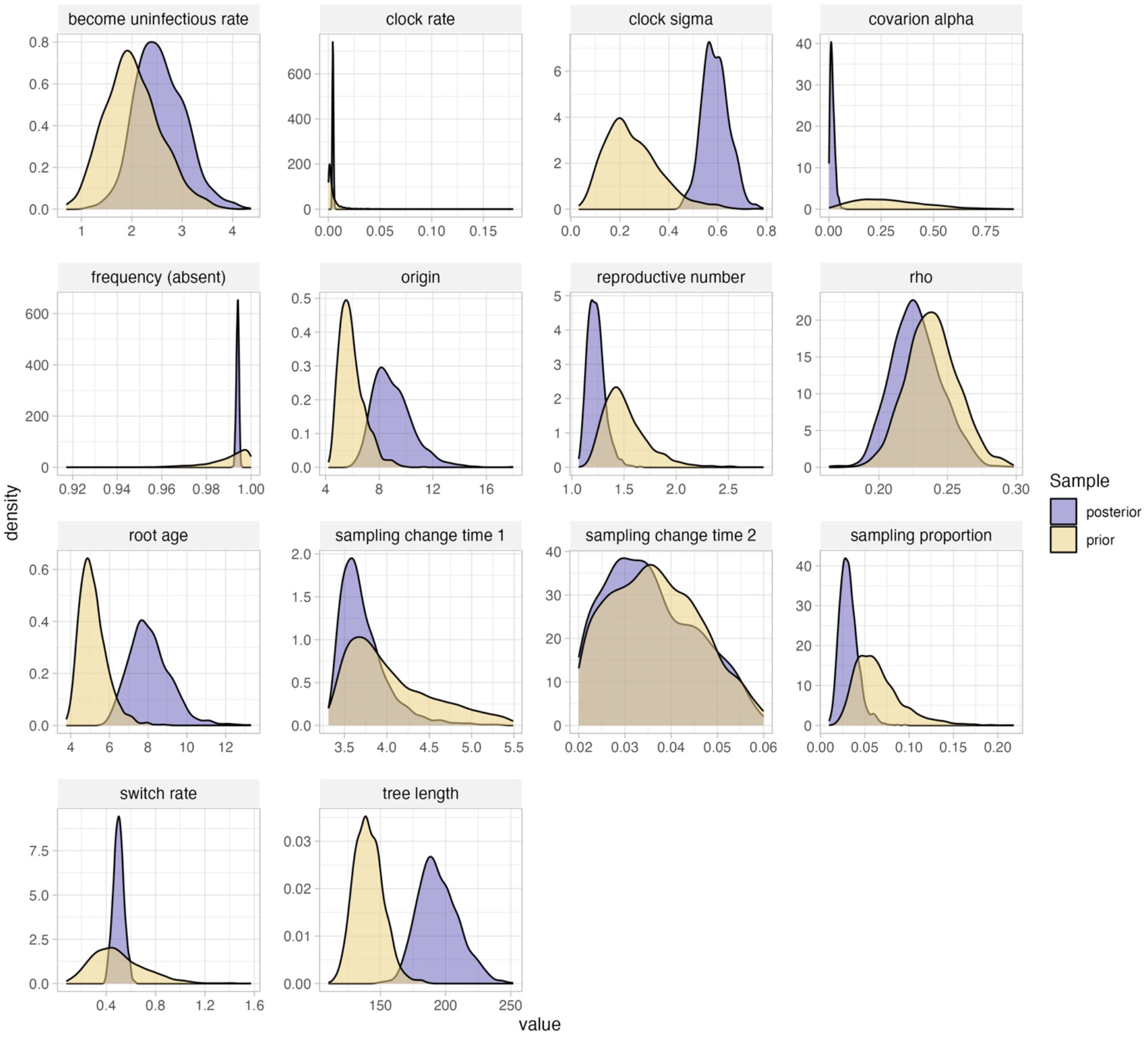
Parameter estimates for a Bayesian tip-dated analysis of a subset of the Indo-European language dataset. Comparison of sampling from the prior (without data) and posterior (with data) shows that the majority of parameters are shifted far from their prior distributions.

**Figure 2.**
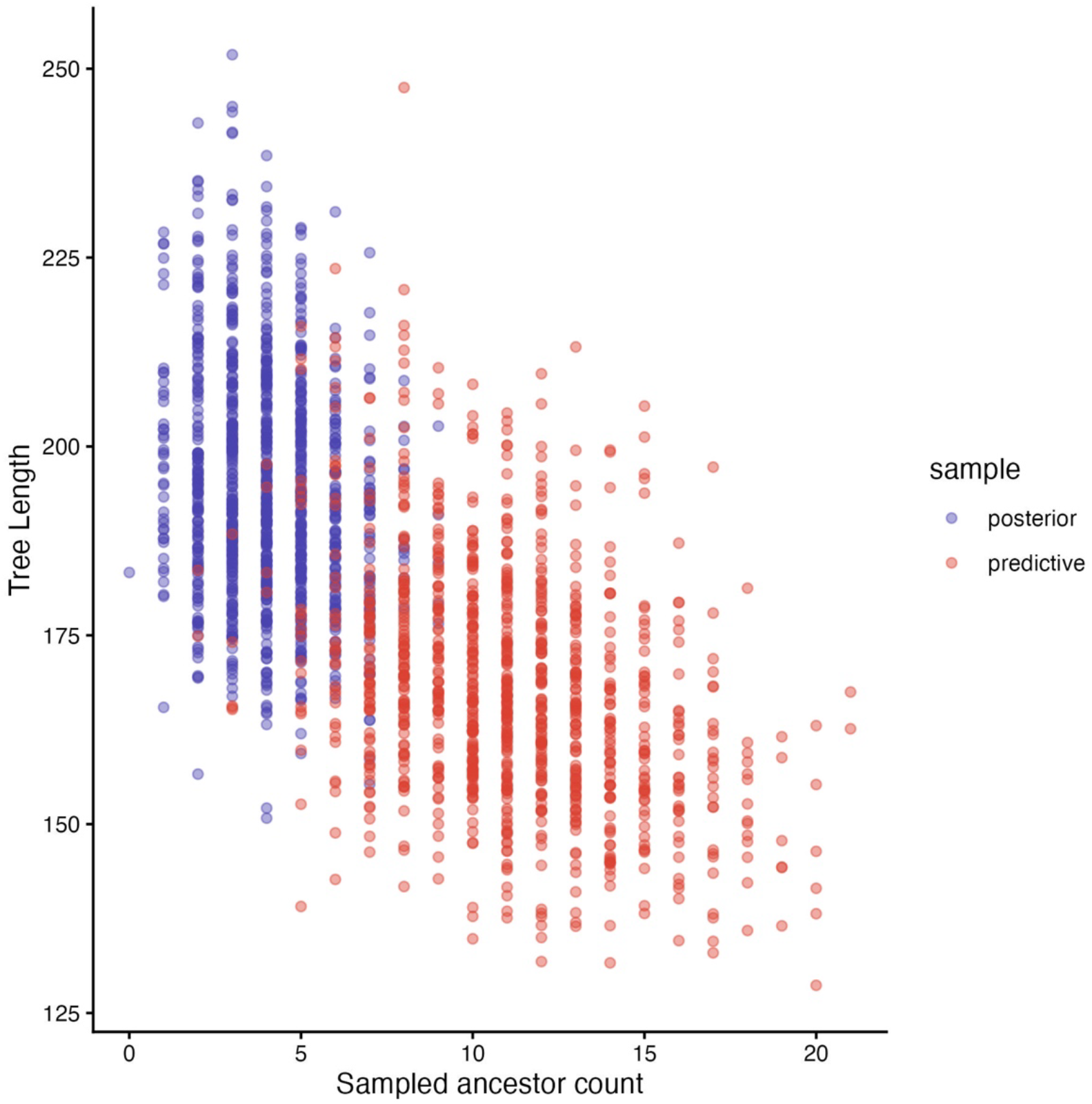
Model misspecification of the tree prior for the Bayesian phylogenetic analysis of Indo-European languages. The posterior and posterior predictive trees diverge slightly on tree length and sampled ancestor count.

Prior SBC of the Indo-European dataset shows good calibration across all parameters, as indicated on the ECDF difference plots (extended data figure 1). The coverage plot (Figure 3) shows good recovery of the parameters of the branch rate model (clock rate and standard deviation) and substitution model (frequencies and covarion alpha parameter). The covarion switch rate is poorly identifiable, but remains well-calibrated, as the estimated distributions deviate little from the prior. Likewise, the parameters of the tree model (origin, reproductive number, sampling proportion, become uninfectious rate, sampling rate change times) are poorly identifiable but well-calibrated. Root age and tree length are well-calibrated and identifiable, as are all of the node dates in the tree (extended data figures 2 and 3).

**Figure 3.**
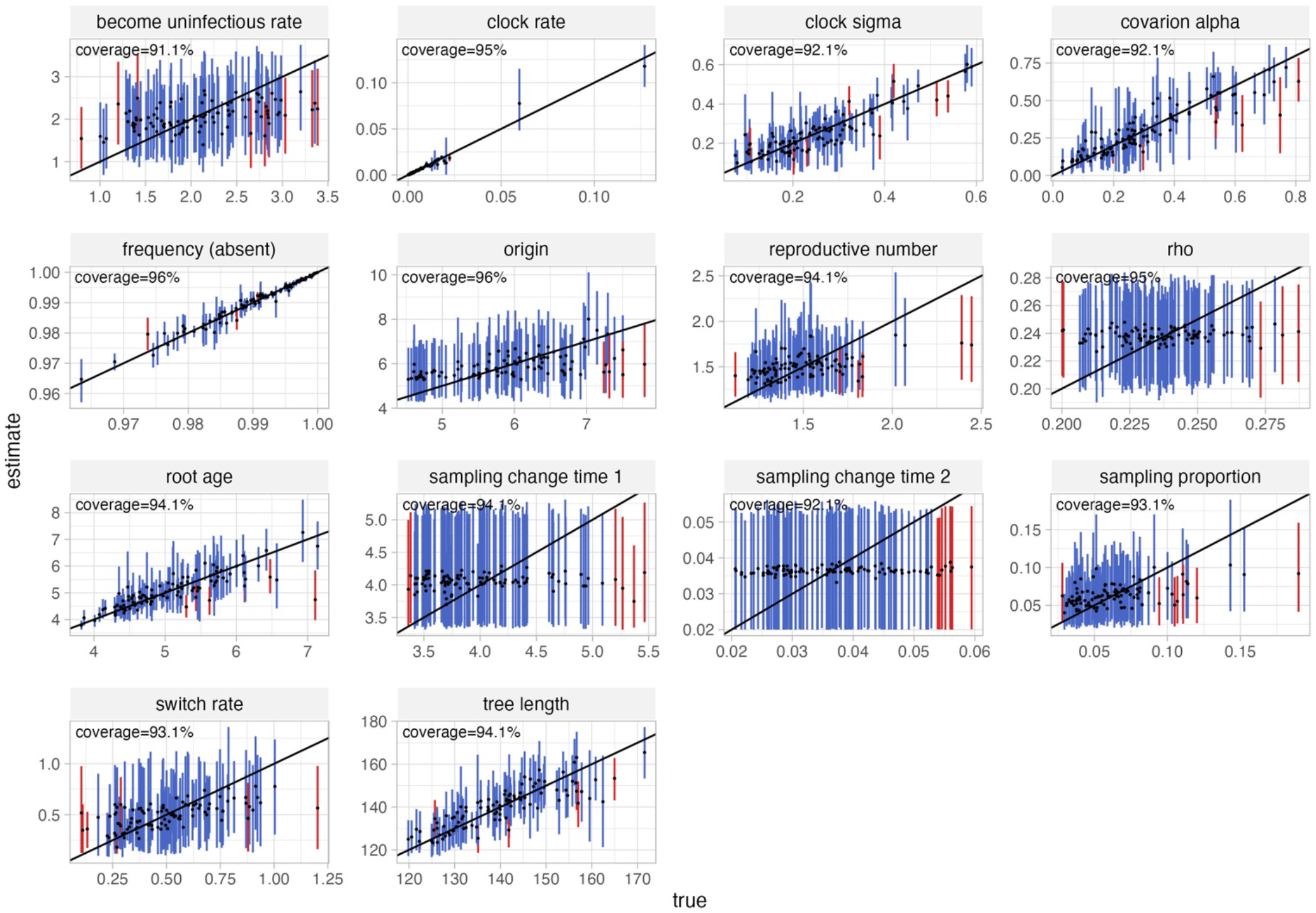
ECDF difference plots of the parameters of the Indo-European tip-dated analysis under posterior SBC. A perfectly calibrated model would show a horizontal line at 0. Deviations that remain within the 95% confidence envelope (light blue) are not considered significant. Estimates for all parameters show good calibration.

Posterior SBC of the Indo-European dataset shows good calibration across all parameters on the ECDF difference plots (Figure 4). However, with the exception of the frequency parameter, there is no further increase in precision for any parameter (Figure 5). Sampling from the augmented posterior results in distributions virtually indistinguishable from the posterior. Most notably, the root age of the tree is estimated at around 8ka in all augmented posteriors, despite the posterior predictive datasets from the tails of the distribution being simulated on trees as young as 6ka or as old as 13ka. This lack of identifiability around the posterior is repeated across all nodes of the phylogeny (extended data figure 4). Nevertheless, the estimates remain well-calibrated since each augmented posterior distribution recapitulates the posterior (extended data figure 5).

**Figure 4.**
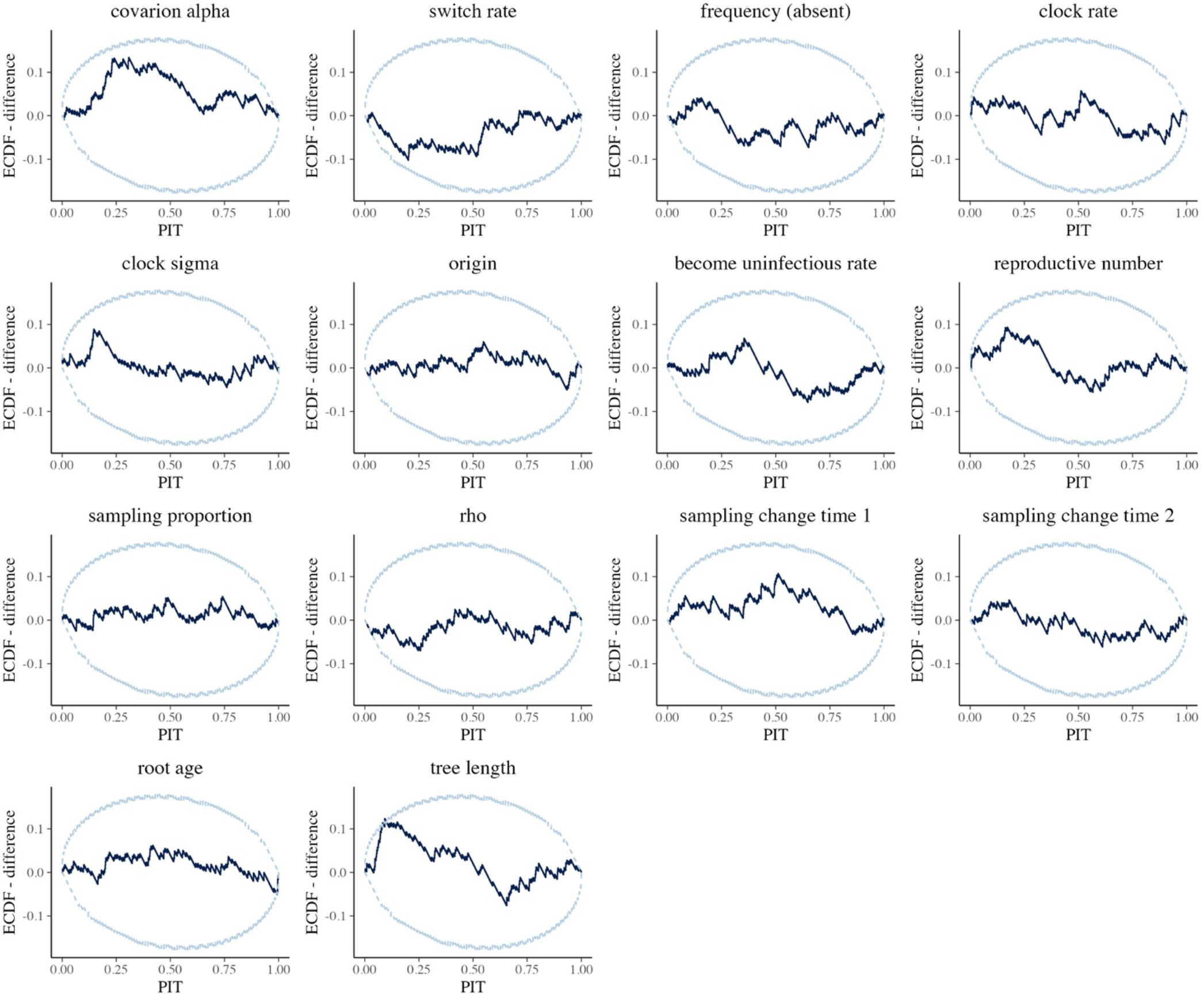
Coverage plot of the parameters of the Indo-European tip-dated analysis under prior SBC. Panels show 95% highest posterior density (HPD) intervals (y axis) against true values (x axis. The diagonal lines represent x=y. HPD intervals are blue when the true value is within the HPD, otherwise red. Coverage refers to the percentage of HPDs which contain the true value, for a well-calibrated analysis this is approximately 95%.

**Figure 5.**
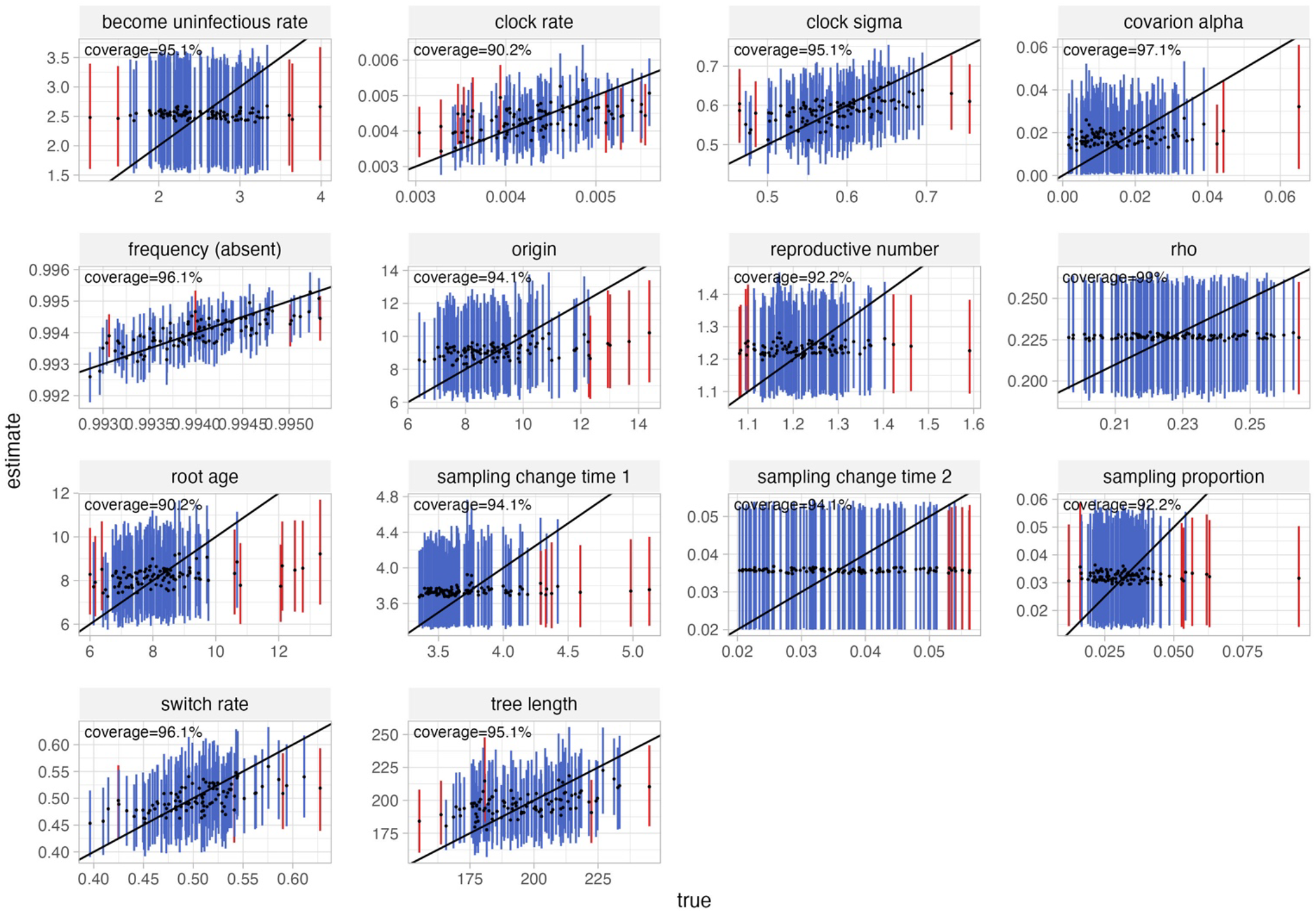
Coverage plot using 95% HPDs of the parameters of the Indo-European tip-dated analysis under posterior SBC. For most parameters the augmented posterior distributions recapitulate the original posterior, with no increase in precision.

Posterior SBC produces very similar results when a simulated dataset is used in place of the empirical data. The coverage plot shows that the distributions for all free parameters, with the exception of the frequency parameter, recapitulate the posterior with no further increase in precision (Figure 6). This holds also for the age distributions of all internal nodes (extended data figure 7). Nevertheless, all parameter and node estimates are well-calibrated (extended data figures 6 and 8).

**Figure 6.**
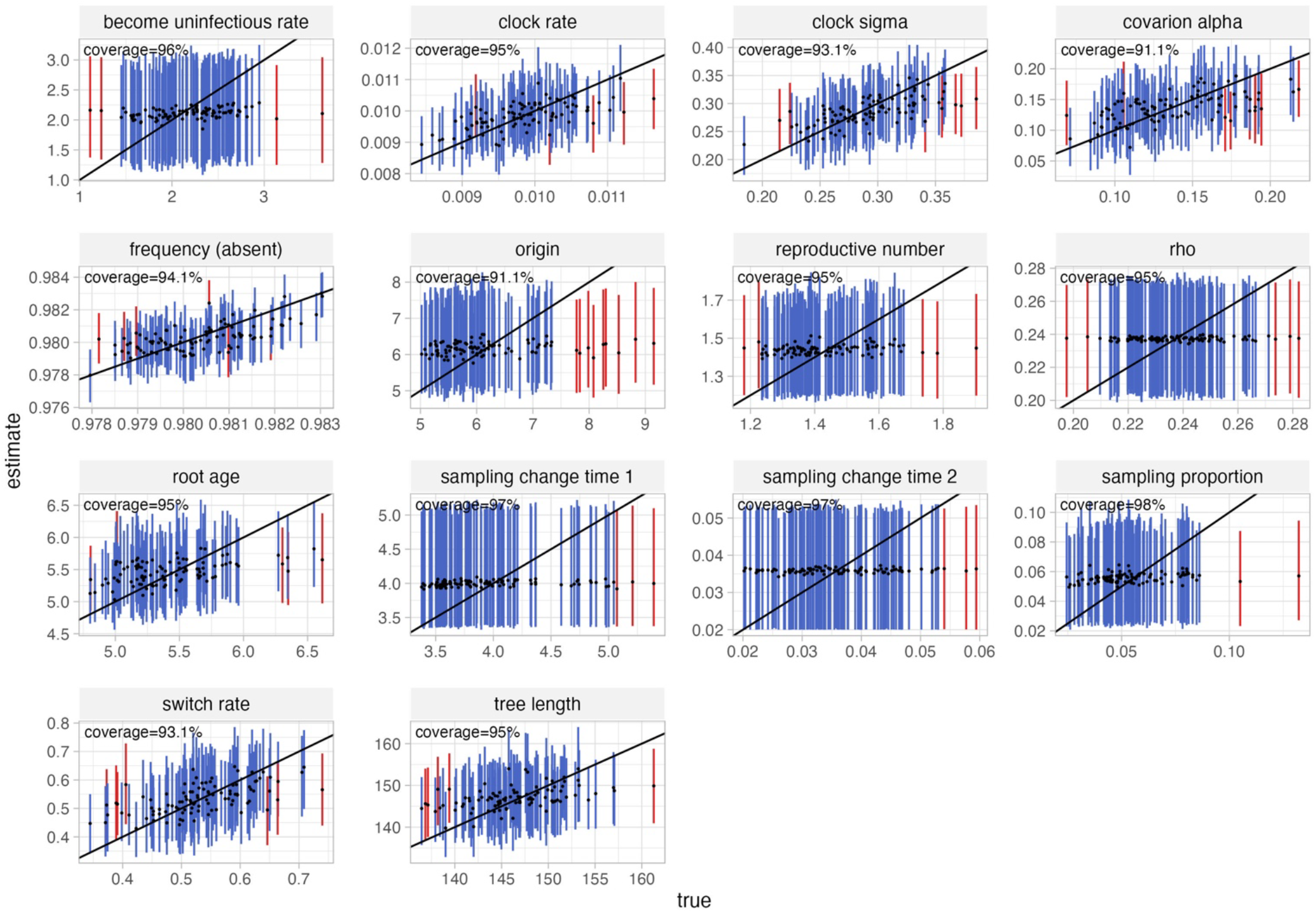
Coverage plot for posterior SBC using a simulated dataset in place of the empirical data. As for the posterior SBC on real data, there is no increase in precision for the estimates of most parameters in the augmented posteriors compared to the posteriors, including the root age. This suggests that this is caused by fundamental constraints on identifiability rather than properties of the empirical data.

### Node-Dated Tabanidae dataset

The majority of parameter estimates in the posterior distribution depart clearly from their effective prior distributions (Figure 7). Misspecification of the tree model is demonstrated by separation of the posterior predictive and posterior trees in the external:internal branch length ratio and gamma metrics (Figure 8).

**Figure 7.**
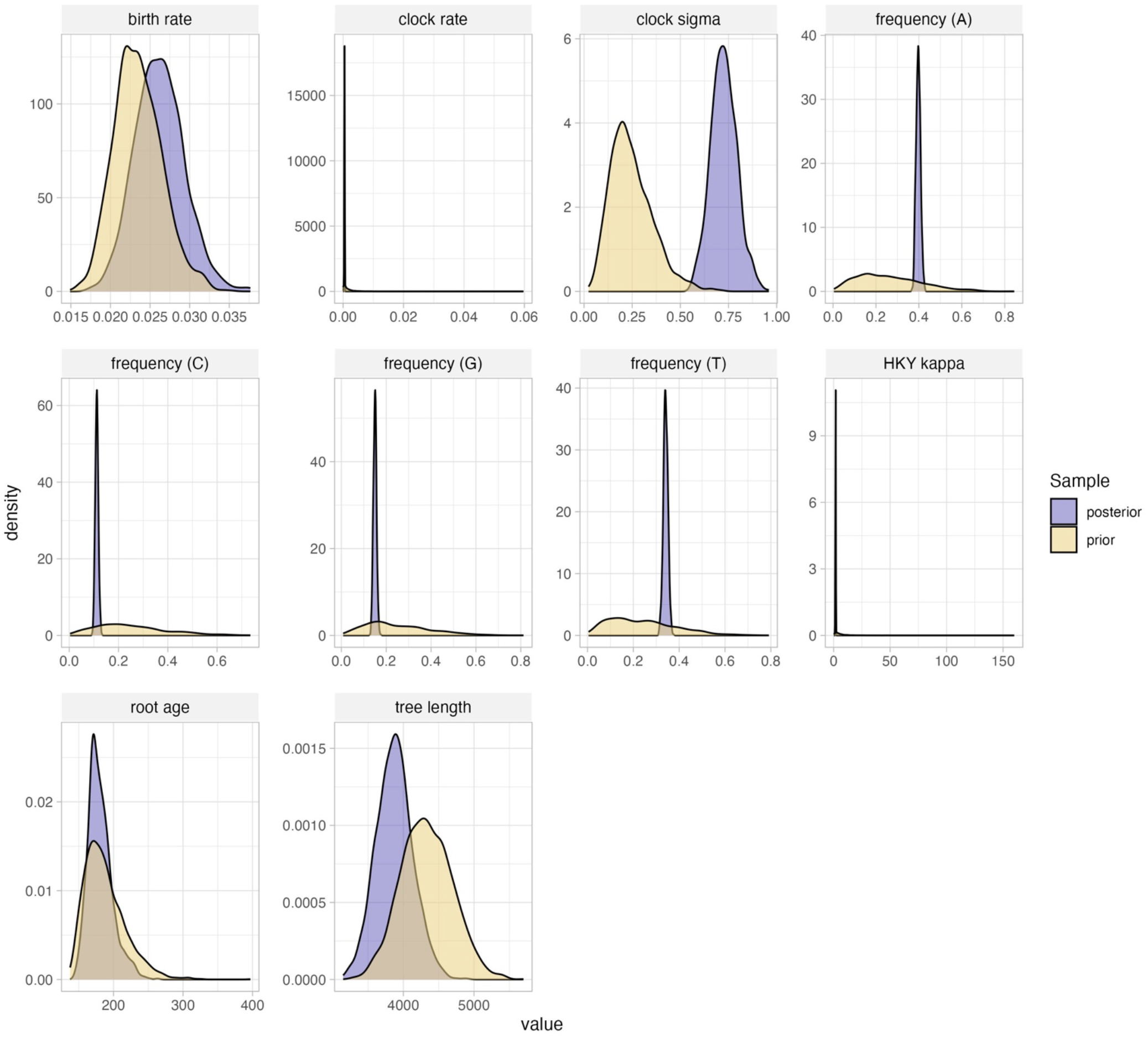
Parameter estimates for a Bayesian analysis of the non-stem 28s sites from the Tabanidae dataset. Comparison of sampling from the prior (without data) and posterior (with data) shows that the majority of parameters are shifted far from their prior distributions.

**Figure 8.**
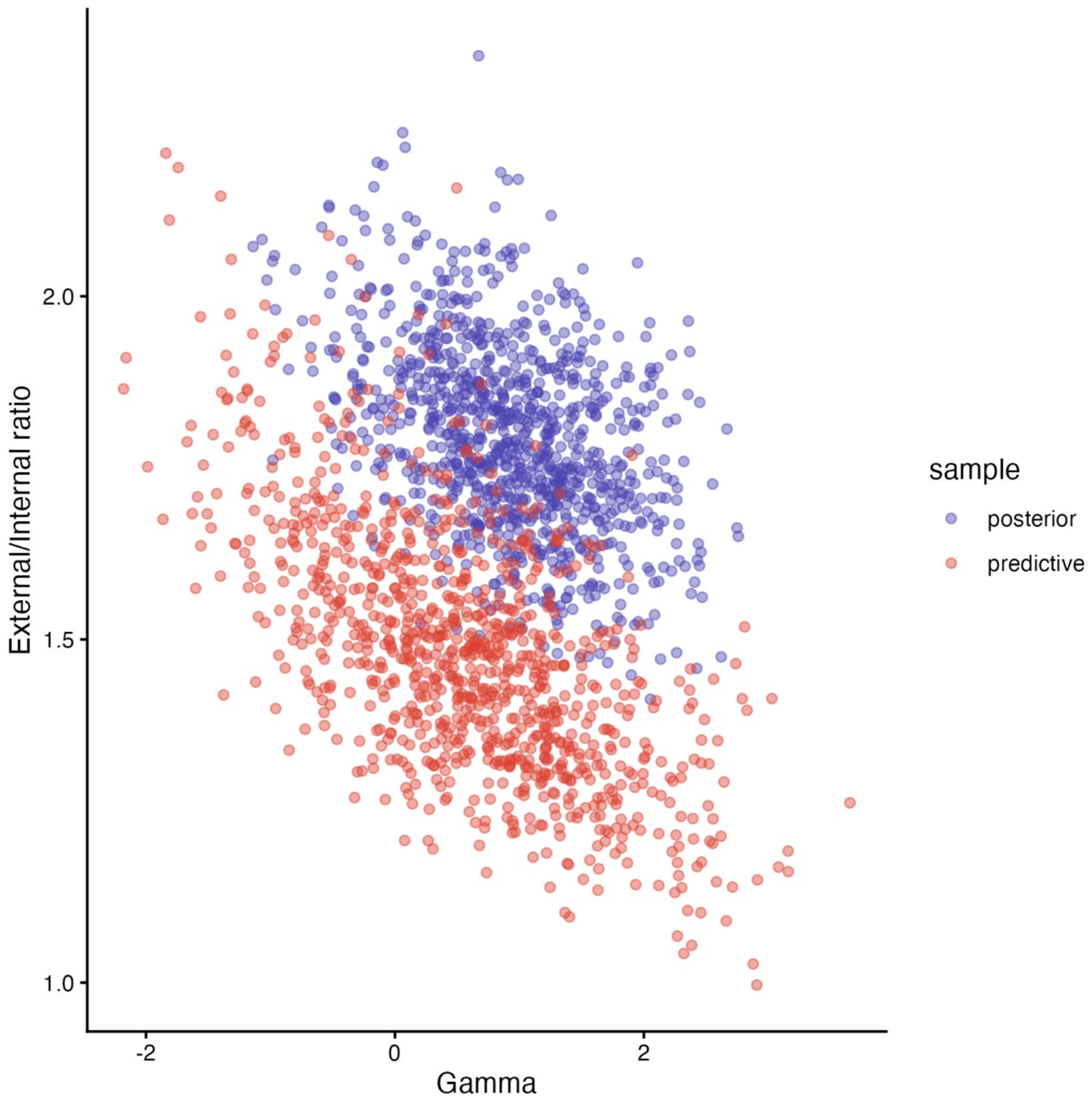
Model misspecification of the tree prior for the Bayesian phylogenetic analysis of Tabanidae. The posterior and posterior predictive trees diverge slightly on external:internal branch length ratio and the gamma statistic.

Under Prior SBC on the Tabanidae dataset, parameters of the substitution model (frequencies, kappa) and branch rate model (clock rate, standard deviation) are well-calibrated and strongly identifiable (extended data figures 9 and 10). The birth rate of the yule model and the root age of the tree are likewise well-calibrated, and weakly identifiable. Age estimates for all nodes are well-calibrated and identifiable, with the exception of the three nodes with calibration densities, for which the distributions follow the prior (extended data figures 11 and 12).

Posterior SBC of the Tabanidae dataset reveals similar results to the tip-dated language dataset above (Figures 9 and 10). There is no increase in precision for the estimates of the root age, tree length, birth rate, clock rate or clock standard deviation. All of the augmented posterior distributions resemble the posterior distribution closely. In contrast, the parameters of the substitution model (frequencies, kappa) do show a further increase in precision in the augmented posteriors. As for the Indo-European dataset, the age estimates for the internal nodes in the posterior SBC test show good calibration but recapitulate the posterior distributions with no increase in precision (extended data figures 13 and 14).

**Figure 9.**
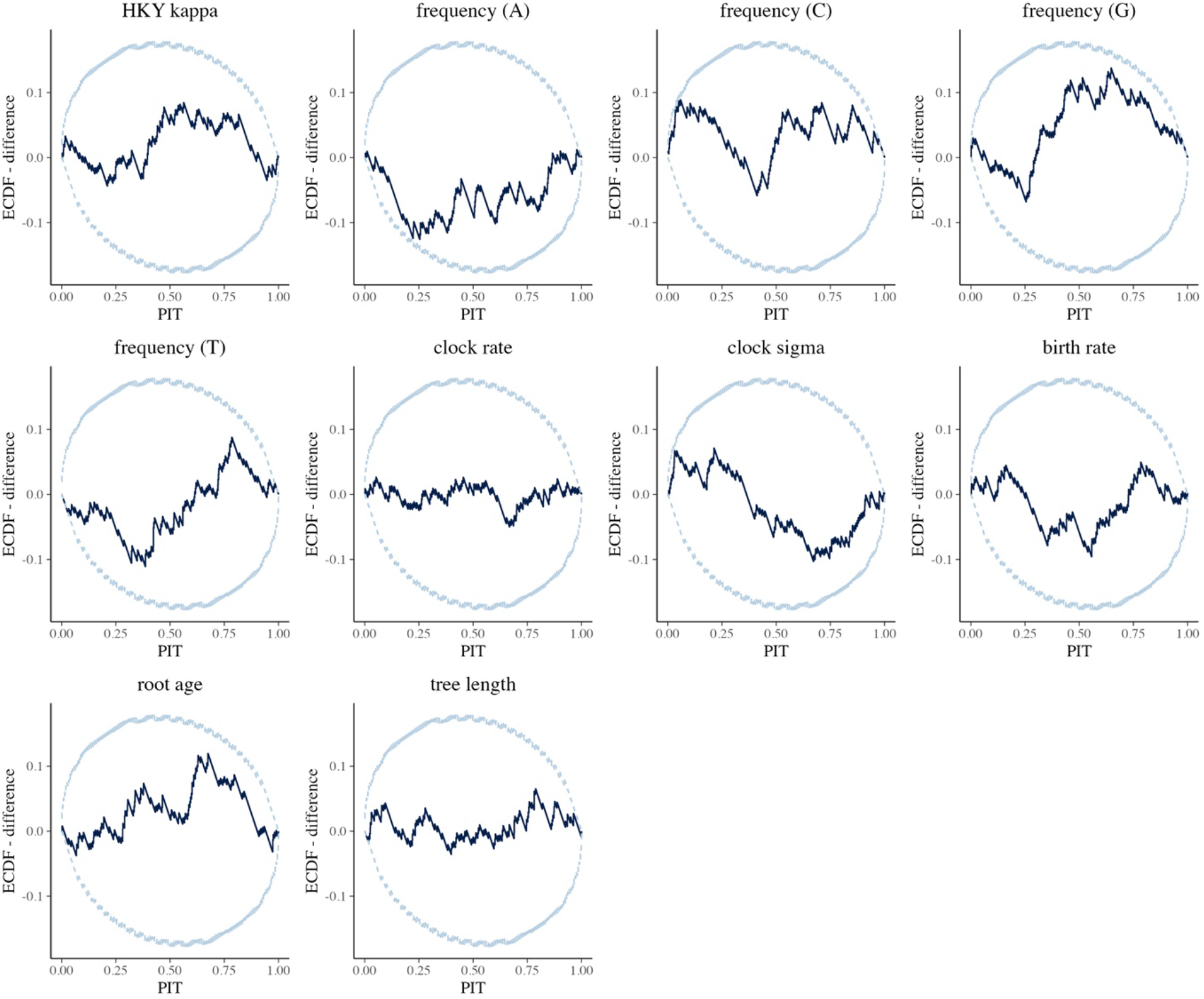
ECDF difference plots of the parameters of the Tabanidae node-dated analysis under posterior SBC. A perfectly calibrated model would show a horizontal line at 0. Deviations that remain within the 95% confidence envelope (light blue) are not considered significant. Estimates for all parameters show good calibration.

**Figure 10.**
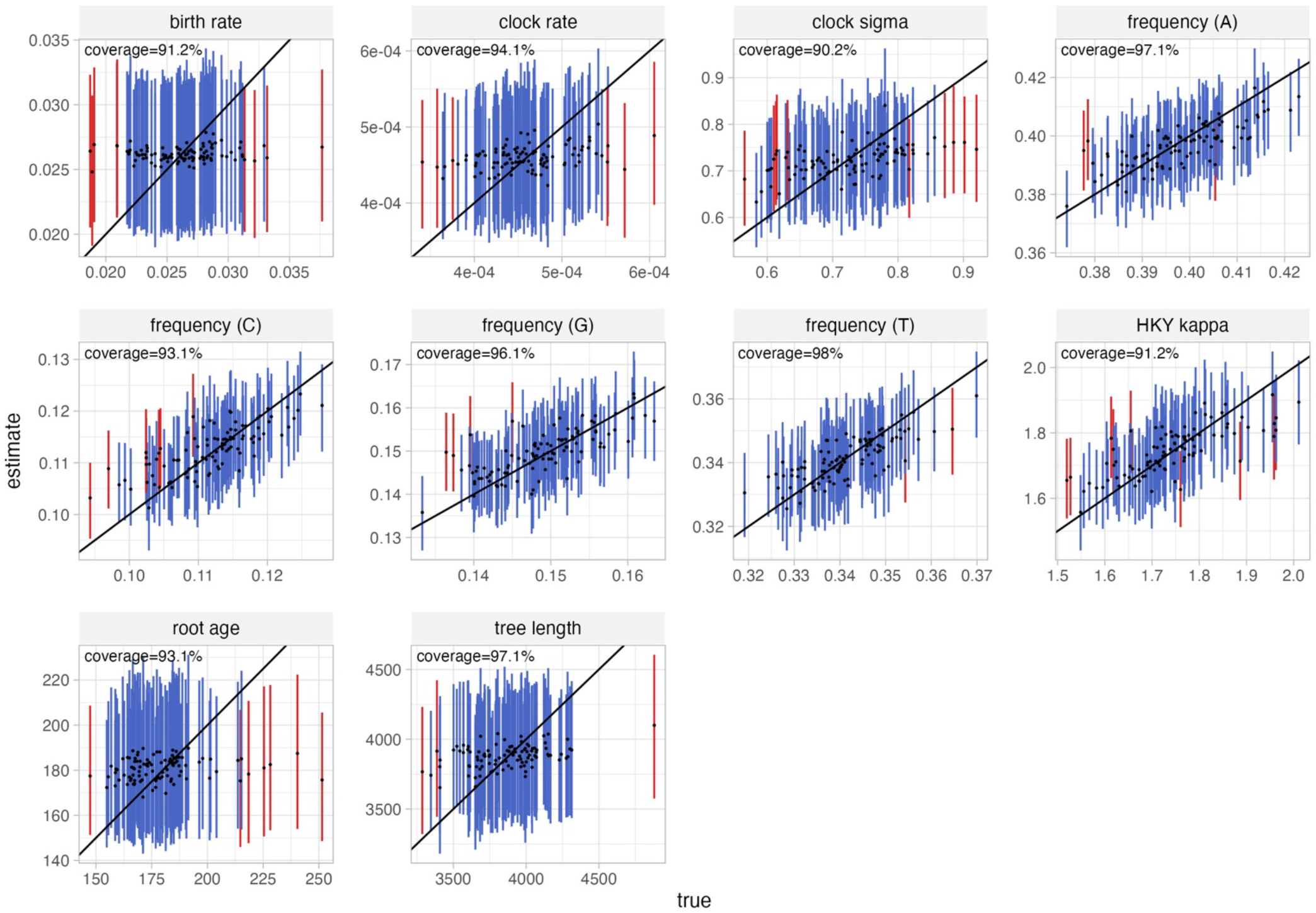
Coverage plot of the parameters of the Tabanidae node-dated analysis under posterior SBC. For most parameters the augmented posterior distributions recapitulate the original posterior, with no increase in precision.

## Discussion

This study is the first to use posterior simulation-based calibration to validate phylogenetic dating methods. It is also the first published study to test node-dating analyses with multiple calibrations using prior SBC. The tests not only validate the inference machinery on empirical data, but show that even in the presence of model misspecification of the tree prior, analyses still produce well-calibrated posteriors. This is a strong indicator that the inference machinery is working as expected. Given the controversy that often surrounds the results of phylogenetic dating analyses, for example debates concerning the ancient origin estimated for the Indo-European language family (Anthony and Ringe 2015, Kassian and Starostin 2025), this is an important validation. This study demonstrates that previous results were not simply the product of software bugs.

The most striking aspect of the results is the lack of any increase in precision for node age estimates under posterior SBC when compared to estimates from the posterior. This lack of increase in precision does not seem to be caused by model misspecification or come characteristic of the empirical data, because when posterior SBC was performed with a simulated prior predictive dataset in place of the empirical data, it produced identical results. This suggests the lack precision is a consequence of fundamental limits, aligning with previous work showing that node-calibrated analyses reach a theoretical limit of precision, even when an infinite number sites are available for analysis (Yang and Rannala 2006; Rannala and Yang 2007; Dos Reis and Yang 2013).

Nevertheless, it is counter-intuitive that when posterior predictive datasets are simulated on either very young or very old trees from the posterior, the augmented posterior distributions do not shift away from the posterior distribution at all: the age of young trees is overestimated and the age of older trees underestimated. This is easily explained when we consider that the data informs branch lengths in terms of substitutions, rather than the time trees themselves. In a hypothetical situation with infinite data, the branch lengths (in terms of expected substitutions per site) are known for certain, but uncertainty in the estimation of node ages remains due to uncertainty in the node or tip calibrations and the relaxed clock branch rates (Yang and Rannala 2006). However, the time trees from the resulting distribution retain identical underlying branch lengths. Therefore, simulating data on any of the resulting trees, including the very youngest or oldest, is directly equivalent and will not change the distribution of node ages in trees estimated from the simulated data.

One caveat to this study is that using MCMC to produce tree samples from the prior introduces some degree of circularity. The same MCMC algorithm is used to sample from the prior and to perform inference. Ideally, trees should instead be produced by direct simulation, independent from the inference machinery being tested (Mendes *et al*. 2025). However, direct simulation is not currently an option for node dating with multiple calibrations. For tip-dated analyses, direct simulation for the birth-death skyline model is available in the R package TreeSim (Stadler 2019), but does not allow a tree model directly equivalent to the Indo-European analysis to be specified. Indeed, direct simulation of the Indo-European tree model is precluded, since it requires a fixed set of extinct taxa with tip date prior distributions. Direct simulation of trees from the tree model parameters produces varying numbers of extant and fossil taxa, which have exact dates. The number of extant taxa is an outcome of the diversification rate parameters (R0, become uninfectious rate) and the fossil taxa are an outcome of diversification and sampling parameters. In one way the taxa and their tip dates more closely resemble data than parameters, and the “prior” in a phylogenetic analysis resembles a posterior distribution, conditional on the observed taxa and their dates. Since direct simulation was not available, tree samples were produced using MCMC for this study.

Clearly, the development of direct simulators for the ever-increasing array of complex phylogenetic tree models is an area in which much future work could be done. Another avenue to explore is the use of different phylogenetic software to generate draws from the prior and to perform inference, for example prior draws could be generated using MCMC in RevBayes (Höhna *et al*. 2016), and then inference performed in BEAST 2. This removes some of the circularity associated with using MCMC to produce samples from the model, and allows distinct software implementations to cross-verify each other.

## Data availability

All data and code for this study, as well as extended figures, are available at figshare under a CC BY 4.0 licence. https://doi.org/10.6084/m9.figshare.31901986

## Competing interests

No competing interests were disclosed.

## Grant information

This work was supported by the Max Plank Society.

## Acknowledgements

I thank Alexei Drummond and Tim Vaughan for comments on an earlier manuscript which inspired this study.

## Notes

https://doi.org/10.6084/m9.figshare.31901986

### Competing Interest Statement

The authors have declared no competing interest.

## References

Anderson, C., Scarborough, M., Jocz, L., Kümmel, M.J., Jügel, T., Irslinger, B., Pooth, R., Liljegren, H., Strand, R.F., Haig, G. and Geupel, U., 2025. The Indo-European Cognate Relationships dataset. Scientific Data, 12(1), p.1541.

Anthony, D.W. and Ringe, D., 2015. The Indo-European homeland from linguistic and archaeological perspectives. Annu. Rev. Linguist., 1(1), pp.199–219.

Berling, L., Klawitter, J., Bouckaert, R., Xie, D., Gavryushkin, A. and Drummond, A.J., 2025. Accurate Bayesian phylogenetic point estimation using a tree distribution parameterized by clade probabilities. PLoS computational biology, 21(2), p.e1012789.

Bouckaert, R., Vaughan, T.G., Barido-Sottani, J., Duchêne, S., Fourment, M., Gavryushkina, A., Heled, J., Jones, G., Kühnert, D., De Maio, N. and Matschiner, M., 2019. BEAST 2.5: An advanced software platform for Bayesian evolutionary analysis. PLoS computational biology, 15(4), p.e1006650.

Cook, S.R., Gelman, A. and Rubin, D.B., 2006. Validation of software for Bayesian models using posterior quantiles. Journal of Computational and Graphical Statistics, 15(3), pp.675–692.

Dawid, A.P., 1982. The well-calibrated Bayesian. Journal of the American statistical Association, 77(379), pp.605–610.

Dos Reis, M. and Yang, Z., 2013. The unbearable uncertainty of Bayesian divergence time estimation. Journal of Systematics and Evolution, 51(1), pp.30–43.

Douglas, J., Zhang, R. and Bouckaert, R., 2021. Adaptive dating and fast proposals: Revisiting the phylogenetic relaxed clock model. PLoS computational biology, 17(2), p.e1008322.

Drummond, A.J., Ho, S.Y.W., Phillips, M.J. and Rambaut, A., 2006. Relaxed phylogenetics and dating with confidence. PLoS biology, 4(5), p.e88.

Drummond, A.J. and Suchard, M.A., 2008. Fully Bayesian tests of neutrality using genealogical summary statistics. BMC genetics, 9(1), p.68.

Duchêne, D.A., Duchêne, S., Holmes, E.C. and Ho, S.Y., 2015. Evaluating the adequacy of molecular clock models using posterior predictive simulations. Molecular Biology and Evolution, 32(11), pp.2986–2995.

Duchêne, S., Bouckaert, R., Duchêne, D.A., Stadler, T. and Drummond, A.J., 2019. Phylodynamic model adequacy using posterior predictive simulations. Systematic biology, 68(2), pp.358–364.

Gabry, J., Simpson, D., Vehtari, A., Betancourt, M. and Gelman, A., 2019. Visualization in Bayesian workflow. Journal of the Royal Statistical Society Series A: Statistics in Society, 182(2), pp.389–402.

Gavryushkina, A., Welch, D., Stadler, T. and Drummond, A.J., 2014. Bayesian inference of sampled ancestor trees for epidemiology and fossil calibration. PLoS computational biology, 10(12), p.e1003919.

Hasegawa, M., Kishino, H. and Yano, T.A., 1985. Dating of the human-ape splitting by a molecular clock of mitochondrial DNA. Journal of molecular evolution, 22(2), pp.160–174.

Höhna, S., Landis, M.J., Heath, T.A., Boussau, B., Lartillot, N., Moore, B.R., Huelsenbeck, J.P. and Ronquist, F., 2016. RevBayes: Bayesian phylogenetic inference using graphical models and an interactive model-specification language. Systematic biology, 65(4), pp.726–736.

Kassian, A.S. and Starostin, G., 2025. Do ‘language trees with sampled ancestors’ really support a ‘hybrid model’for the origin of Indo-European? Thoughts on the most recent attempt at yet another IE phylogeny. Humanities and Social Sciences Communications, 12(1), pp.1–10.

Mahani, A.S., Hasan, A., Jiang, M. and Sharabiani, M.T., 2016. Stochastic Newton Sampler: The R Package sns. Journal of Statistical Software, 74, pp.1–33.

Mendes, F.K., Bouckaert, R., Carvalho, L.M. and Drummond, A.J., 2025. How to validate a Bayesian evolutionary model. Systematic Biology, 74(1), pp.158–175.

Morita, S.I., Bayless, K.M., Yeates, D.K. and Wiegmann, B.M., 2016. Molecular phylogeny of the horse flies: a framework for renewing tabanid taxonomy. Systematic entomology, 41(1), pp.56–72.

Penny, D., McComish, B.J., Charleston, M.A. and Hendy, M.D., 2001. Mathematical elegance with biochemical realism: the covarion model of molecular evolution. Journal of Molecular Evolution, 53(6), pp.711–723.

Pybus, O.G. and Harvey, P.H., 2000. Testing macro–evolutionary models using incomplete molecular phylogenies. Proceedings of the Royal Society of London. Series B: Biological Sciences, 267(1459), pp.2267–2272.

Rannala, B. and Yang, Z., 2007. Inferring speciation times under an episodic molecular clock. Systematic biology, 56(3), pp.453–466.

Rambaut, A., Drummond, A.J., Xie, D., Baele, G. and Suchard, M.A., 2018. Posterior summarization in Bayesian phylogenetics using Tracer 1.7. Systematic biology, 67(5), pp.901–904.

Säilynoja, T., Bürkner, P.C. and Vehtari, A., 2022. Graphical test for discrete uniformity and its applications in goodness-of-fit evaluation and multiple sample comparison. Statistics and Computing, 32(2), p.32.

Säilynoja, T., Schmitt, M., Bürkner, P.C. and Vehtari, A., 2026. Posterior SBC: simulation-based calibration checking conditional on data. Statistics and Computing, 36(2), p.78.

Stadler, Tanja (2019). TreeSim: Simulating Phylogenetic Trees. R package version 2.4. url: https://CRAN.R-project.org/package=TreeSim.

Stadler, T., Kühnert, D., Bonhoeffer, S. and Drummond, A.J., 2013. Birth–death skyline plot reveals temporal changes of epidemic spread in HIV and hepatitis C virus (HCV). Proceedings of the National Academy of Sciences, 110(1), pp.228–233.

Talts, S., Betancourt, M., Simpson, D., Vehtari, A. and Gelman, A., 2018. Validating Bayesian inference algorithms with simulation-based calibration. arXiv preprint arXiv:1804.06788.

Tuffley, C. and Steel, M., 1998. Modeling the covarion hypothesis of nucleotide substitution. Mathematical biosciences, 147(1), pp.63–91.

Yang, Z. and Rannala, B., 2006. Bayesian estimation of species divergence times under a molecular clock using multiple fossil calibrations with soft bounds. Molecular biology and evolution, 23(1), pp.212–226.

Zhang, R. and Drummond, A., 2020. Improving the performance of Bayesian phylogenetic inference under relaxed clock models. BMC evolutionary biology, 20(1), p.54.

